# Tandem bromodomains of BRD4 cooperatively read poly-acetylated nucleosomes to enhance chromatin engagement and regulate breast cancer phenotypes

**DOI:** 10.64898/2026.04.22.719657

**Authors:** Nathaniel T Burkholder, Shwu-Yuan Wu, Jackson Handy, Cole Bertram, Matthew R. Marunde, Irina K. Popova, Krzysztof Krajewski, Cheng-Ming Chiang, Brian D Strahl

## Abstract

Bromodomain-containing protein 4 (BRD4) is an acetyl-lysine reader protein implicated in transcriptional control and oncogenesis, yet how its tandem bromodomains (BD1-2) contribute to nucleosomal engagement remains unresolved. Here we show that the tandem bromodomains of BRD4 cooperatively engage poly-acetylated histone H4 tails and nucleosomes *in vitro* and promote chromatin association in human cells. In stringent peptide pull-down and nucleosome-based biolayer interferometry assays, isolated BRD4 bromodomains bind weakly to poly-acetylated histone peptides and nucleosomes, whereas the tandem BD1-2 module binds much more robustly. These results closely mirror our observations in mammalian cells, where truncations lacking either bromodomain or pocket-disrupting mutations in either domain reduced chromatin association, with dual pocket disruption causing the strongest defect. In the BRD4 short isoform (BRD4-S), maximal chromatin association additionally required the region C-terminal to the BD2, which contains the basic residue-enriched interaction domain (BID) and extraterminal domain (ET), consistent with a multivalent chromatin engagement mechanism beyond the bromodomains alone. Functionally, dual pocket disruption attenuated BRD4-S-dependent breast cancer phenotypes, including impaired growth and reduced transwell migration. Together, these findings define how tandem bromodomains and adjacent BRD4-S regions cooperate to stabilize chromatin residence and inform therapeutic strategies aimed at more precisely disrupting BRD4 function.

**Graphical Abstract:** 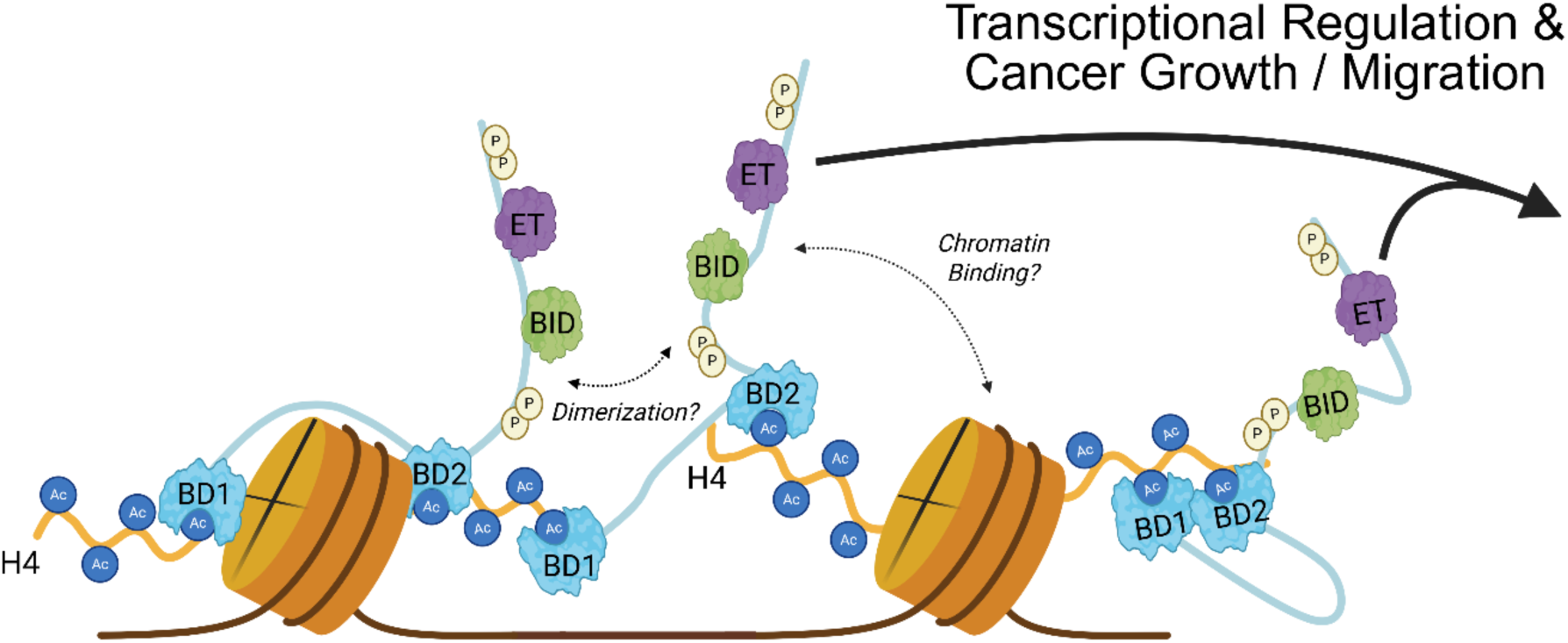

## Introduction

Dysregulation of epigenetic mechanisms can impact many essential cellular functions including transcription, DNA repair, and DNA replication, and is increasingly recognized as a hallmark of many cancers [1–6]. The histone code hypothesis proposes that specific combinations of histone post-translational modifications (PTMs) orchestrate complex regulatory outcomes through interactions with epigenetic reader proteins [7,8]. These readers interpret combinatorial histone marks, dictating diverse biological processes in both normal physiology and disease [9,10]. Consequently, targeting epigenetic reader proteins, particularly those recognizing histone acetylation, has emerged as a promising therapeutic strategy, exemplified by ongoing clinical trials of bromodomain and extraterminal (BET) family inhibitors [11,12].

BET inhibitors primarily target the bromodomains of BRD2, BRD3, BRD4, and BRDT, with BRD4 notably implicated in multiple malignancies that include ovarian, lung, and breast cancer [13,14]. BRD4 functions as a critical epigenetic reader that can engage acetylated chromatin and promote transcription initiation and elongation [15,16], in part through association with sequence-specific transcription factors [17] and RNA polymerase II pause-release mechanisms [18]. In addition to being ubiquitously expressed, BRD4 regulates a broad range of genes important for normal lineage determination and disease development [19,20]. Given this, BRD4 has emerged as a major drug target for several cancers, with specific and potent bromodomain inhibitors capable of rapidly perturbing its reader function and downstream transcriptional programs [21,22].

Intriguingly, BRD4 has two different ubiquitously expressed isoforms: full-length BRD4-L (long), and BRD4-S (short, Fig. 1A) which lacks C-terminal domains involved in coordinating elongation complexes that mediate promoter-proximal pause-release at transcription start sites [18]. Studies of BRD4-L and BRD4-S have identified distinct functions between the isoforms, most notably in disease. For example, in triple-negative breast cancer (TNBC) BRD4-S is upregulated and promotes oncogenic phenotypes, whereas BRD4-L appears to suppress oncogenesis [23]. These opposing isoform activities highlight that bromodomain targeting of BRD4 may not yield a single desired molecular outcome and motivate further interrogation into how each BRD4 isoform engages chromatin and contributes to cellular phenotypes during inhibitor treatment.

**Figure 1.**
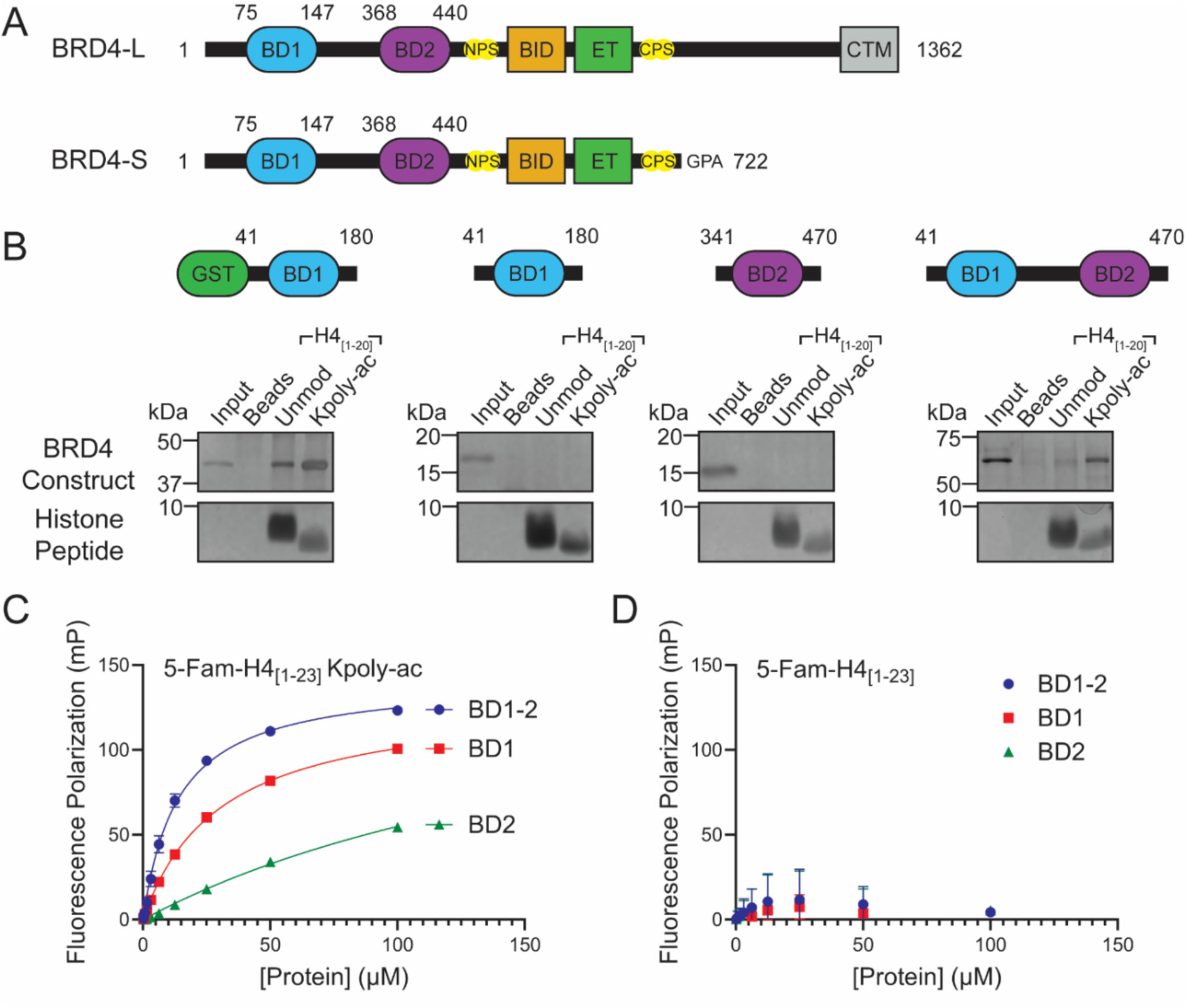
The BRD4 tandem bromodomains stably associate with poly-acetylated histone peptides in pull-down assays. (**A**) Domain diagrams of BRD4-L and BRD4-S isoforms. Indicated are bromodomains 1 and 2 (BD1/2), N- and C-terminal phosphorylation sites (NPS/CPS), basic interaction domain (BID), extraterminal (ET) domain, and carboxy-terminal motif (CTM). (**B**) Silver-stained gel of histone H4_[1–20]_ peptide pull-down (unmodified or K5acK8acK12acK16ac) pull-downs using the indicated BRD4 bromodomain constructs under stringent wash conditions. Differences in gel migration and signal intensity between H4 peptides likely reflect their lysine acetylation status. Pull-down experiments were performed in triplicate. (**C-D**) Fluorescence polarization assays using the indicated BRD4 constructs and either (**C**) H4_[1–23]_K5acK8acK12acK16acK20ac or (**D**) H4_[1–23]_ peptides labeled with 5-FAM. Experiments were conducted in triplicate and the binding curves were fitted using one-site nonlinear regression (Prism, Graphpad).

BRD4 harbors two conserved bromodomains (BD1 and BD2) separated by a flexible linker [15,24]. Both domains recognize poly-acetylated H3 and H4 tails in the context of peptides [16,25,26] and predominantly poly-acetylated H4 nucleosomes albeit with some synergy towards nucleosomes with both H3 and H4 poly-acetylation [27]. Notably, BD1 possesses much higher affinity towards poly-acetylated histones than BD2 [25]. However, studies still point to an important role for BD2 given the tandem BD1-2 module binds more strongly than isolated BD1 to poly-acetylated nucleosomes [27]. Additionally, mutation of the acetyl-lysine binding pocket of either BD1 or BD2 impairs BRD4 chromatin association in cells after photobleaching to similar levels [26]. Along with other studies demonstrating cooperation of multiple histone reader domains [28–33], these findings suggest that the individual BD1 and BD2 domains alone may inadequately recapitulate BRD4’s observed chromatin-binding properties. Moreover, emerging work suggests BRD4-nucleosomal engagement can also involve contacts beyond acetyl-lysine recognition, underscoring the need to define how BD1 and BD2 contribute within multivalent chromatin binding [34,35].

In this study, we investigated the combinatorial mechanism by which BRD4 bromodomains recognize poly-acetylated histones. This work was inspired by the observation that removing the GST tag capable of dimerization from recombinant BD1 abrogated its binding to acetylated histones in bead-based pull-down experiments. This suggested that combinatorial readout by tandem BD1-2 could be critical for BRD4’s capacity to engage chromatin, despite the well-known lower affinity of BD2 for acetylated histones [25,27]. Using a series of biochemical assays, including peptide pull-downs, fluorescence polarization, and nucleosome-based biolayer interferometry, we show that multivalent engagement by the BRD4 tandem BD1-2 markedly enhances stable association with H4 poly-acetylated tails and nucleosomes, particularly under more stringent assay conditions. Moreover, cellular chromatin association experiments reveal that disruption of either BD1 or BD2 acetyl-binding pocket significantly compromises chromatin association, whereas simultaneous disruption nearly abolishes it. In the context of BRD4-S, mutation of both BD1 and BD2 pockets was required to reduce chromatin association in cells, indicating that regions outside the bromodomains also contribute. Consistent with this, the C-terminal portion of BRD4-S (residues 440-719) was required for near-maximal chromatin engagement, potentially through additional interactions with nucleosomes or other chromatin-associated factors, including other BRD4 molecules. Finally, mutation of both bromodomains in BRD4-S disrupted its ability to complement growth and migration phenotypes in a triple-negative breast cancer model cell line (MDA-MB-231). Together, these findings support a model in which the tandem bromodomains and additional regions in BRD4-S multivalently engage poly-acetylated nucleosomes in chromatin to regulate cellular function in both normal and cancer contexts, with important implications for how BET inhibition perturbs BRD4-dependent gene expression and phenotype.

## Materials and Methods

### Protein expression and purification

Human BRD4 sequences coding amino acids for BD1: 41-180, BD2: 341-470, and BD1-2: 41-470 were cloned into pGEX6P vectors and expressed as GST-linked proteins in *E. coli* BL21(DE3) cells using standard protocols. GST-linked BRD4 bromodomain constructs were extracted in lysis buffer (500 mM NaCl, 50 mM HEPES pH 7.5, 1 g/L lysozyme, 1 μL/10 mL Thermo Pierce Universal Nuclease, 1 mM PMSF, and 1 mM DTT) and purified over Pierce glutathione agarose resin (Thermo) in wash buffer (200 mM NaCl, 25 mM HEPES pH 7.5, and 1 mM DTT). Proteins were eluted in wash buffer supplemented with 10 mM glutathione and re-pH’d to 7.5. These proteins were processed with ∼60 μg/mL GST-precision protease dialyzed O/N in wash buffer, ran back through clean glutathione resin to filter out GST, and further purified over a Superdex 16/600 75 pg size exclusion column (Cytiva) in filtered wash buffer. Elution fractions were evaluated using a size standard curve and gel electrophoresis. Samples containing purified BRD4 bromodomains were collected, concentrated over 10 kDa Amicon ultracentrifugal filters (Millipore) to ∼2-10 mg/mL based on Nanodrop quantification, supplemented with 10% glycerol, and aliquoted and flash frozen in liquid nitrogen prior to storage at -80°C.

### Peptide pull-down assays

Peptide pull-downs were conducted in binding buffer (25 mM HEPES pH 7.5, 150 mM NaCl, 0.1% NP-40). 50 picomoles of the indicated BRD4 bromodomain protein were incubated with the indicated biotinylated peptide in 500 μL of binding buffer, rotating at 4°C overnight. Five μL of washed streptavidin magnetic beads (Pierce) were resuspended in 150 μL binding buffer, added to the sample. The binding mixture was incubated, rotating at 4°C for 1 hour. The beads were washed 3 times with 1 mL of binding buffer and resuspended in SDS loading dye. Proteins bound to the beads were resolved by SDS-page and SilverQuest Silver Staining Kit (Thermo) per manufacturer’s instructions.

### Fluorescence polarization assays

Fluorescence polarization assays were performed under standard buffer conditions (25 mM HEPES pH 7.5, 150 mM NaCl, 0.1% NP-40) in black flat-bottom 384-well plates (Corning). Serial dilutions of BRD4 bromodomain constructs were incubated with indicated 5-FAM-labelled peptides at 80 nM for 60 minutes at room temperature to allow for equilibrium binding. Fluorescence polarization was measured using a CLARIOstar PLUS plate reader (excitation: 482-16 nm; emission: 530-40 nm; gain: 1000, focal distance: 7.3 mm). Polarization values were plotted against protein concentration and curves were fitted using one-site nonlinear regression model in GraphPad Prism.

### Biolayer interferometry

BRD4 bromodomain nucleosome binding kinetics were determined using a ForteBio Octet Red384 instrument on streptavidin-coated (SA) biosensors (Sartorius). In black flat bottom 96 well plates, 2-fold dilution series of either BRD4 BD1-2 or BD1 from 25 – 0.34 μM as well as 12.5 nM unmodified or H4 poly-acetylated nucleosomes ([H4K5acK8acK12acK16ac]_2_, EpiCypher) were prepared in binding buffer (150 mM NaCl, 25 mM HEPES pH 7.5, 0.1% NP-40). Experimental runs consisted of the following steps: 1) buffer baseline for 120s baseline, 2) nucleosome loading to ∼1 nm response (∼300s), 3) buffer wash for 120s, 4) BRD4 association for ∼300s, and 5) BRD4 dissociation in buffer for ∼300s. Experiments were run at 37°C, 10 Hz, and 1000 rpm shaking. Binding curves were fitted to a 2:1 heterogeneous ligand model in Octet Analysis Software (Sartorius).

### Luminex (Captify^TM^) assays

This study uses a nucleosome nomenclature recently devised for accurate scientific communication in the chromatin field [36]. Here any distinguishing histones in a fully-defined semi-synthetic nucleosome are indicated (*e.g.*, as in ([H4K12ac]_2_)), with other positions not denoted should be understood as unmodified major histones.

GST-BRD4 BD1-2 nucleosome binding kinetics were examined using a high sensitivity, flow-based Intelliflex instrument (Luminex). Designer nucleosomes (dNucs; EpiCypher) were made using native chemical ligation to yield full-length ‘scarless’ histones [37,38]. Nucleosome panels were generated as previously [32,39]. In brief, 50 nM of biotinylated nucleosomes were complexed with 2x10^6^ beads/mL of MagPlex-Avidin dyed beads (Luminex) followed by magneting, washing (50 mM Tris pH 7.5, 0.1% Tween-20, 0.5% BSA, 1 mM BME, and 10% glycerol), bead counting (Bio-Rad), and finally combining each in a storage buffer at equi-bead number (10 mM sodium cacodylate 7.5, 0.1% Tween-20, 0.5% BSA, 1 mM BME, and 50% glycerol). Luminex nucleosome panels consisted of the following 601 147 bp DNA-biotinylated nucleosomes (all from EpiCypher): unmodified (rNuc), 16-0006; H4K12 acetylated ([H4K12ac]_2_), 16-0312; H4 poly-acetylated ([H4K5acK8acK12acK16ac]_2_), 16-0313; H3 poly-acetylated ([H3K4acK9acK14acK18ac]_2_), 16-0336; H4 N-terminal tail truncated ([H4ΔN15]_2_), 16-0018; and trypsin-digested tailless, 16-0027. For the 96-well plate assays, GST-BRD4 BD1-2 was titrated at 2x concentration in 75, 150, or 225 mM NaCl buffer (25 mM HEPES pH 7.5, 0.1% BSA, 0.1% Tween-20, 1 mM BME) and mixed 1:1 with 2x nucleosome-bead panel for a 30 min RT incubation shaking at 600 rpm. After incubation, the plates were magneted and washed in buffer before addition of 1:2000 anti-GST (Pocono Farms). After another 30 min incubation, the plates were magneted and washed again before adding 1:200 anti-rabbit-phycoerythrin (PE; Thermo, P-2771MP). After a final 30 min incubation, the plates were magneted and washed before reading in an Intelliflex. Each well was read until 50 of each bead was counted and median PE fluorescence for each bead region detected was determined. Control experiments were conducted in wells with only nucleosome-bead panels added and use of a general histone binding antibody (Millipore, MAB3422) at 1:200 prior to detection with a mouse PE secondary at 1:200 (Thermo, P-852).

### Chromatin association assays

HEK293T cells were transfected with BRD4 bromodomain complexes using Lipofectamine 3000 according to manufacturer’s protocol (ThermoFisher). The cells were then selected in DMEM + 5 μg/mL blasticidin for 3 days. Chromatin association assays were performed as previously described, with slight modification [40]. Cells were harvested by scraping and resuspended in 100 μL CSK buffer (10 mM PIPES pH 7.0, 340 mM sucrose, 100 mM NaCl, 3 mM MgCl_2_, 1x Sigma complete protease inhibitor cocktail, 1 mM DTT) supplemented with 0.1% Triton X-100 and incubated on ice for 10 minutes. 50 μL was combined with 0.5 μL Pierce Universal Nuclease and saved (total fraction). The remaining cell sample was centrifuged at 1300 × g for 5 minutes at 4°C. The supernatant was collected (soluble fraction). The pellet was washed without resuspension with 125 μL of CSK buffer and spun again at 1300 × g for 1 minute at 4°C. The supernatant was discarded and the pellet was resuspended in 50 μL CSK buffer supplemented with 0.1% Triton X-100 and 1:100 Pierce Universal Nuclease. The suspension was incubated on ice for 30 minutes then sonicated with a Sonifier Cell Disruptor 250 (5x 1-second pulses; 40% duty cycle, output control 4) and saved (chromatin fraction). All fractions were resolved by SDS-PAGE and western blotting for detection using the following antibodies: 1:2000 M2 anti-FLAG (Sigma, F3175), 1:10000 anti-H3 Pan (Pocono), 1:1000 anti-α-tubulin (Cell Signaling, 2144).

### AlphaFold modeling and structural visualization

Modeling was performed using the AlphaFold 3 Server (Google). Sequences for the canonical human histones including H4K12 acetylation added, the 147-bp core of the 601 Widom DNA sequence [41], and BRD4-S (Uniprot 060885) were used as inputs for the modeling. The structure of the top AlphaFold model was visualized in ChimeraX (UCSF). Distance constraints between BD2 residues and the nucleosomal phosphodiester backbone were determined using ChimeraX’s structural analysis distance tool.

### Breast cancer viability and transwell migration assays

TNBC cells expressing doxycycline-inducible shBRD4-S [23] were electroporated with PB2-TetOn-3xFLAG-BRD4-S (1-719) expression plasmids with or without phenylalanine (F) substitution for Y37, Y390, or both residues (2xYF). Upon selection with 0.75 μg/mL puromycin and 7.5 μg/mL blasticidin in DMEM + 10% FBS supplemented with 1x plasmocin, primocin, and Pen/Strep, the cells were then treated with 2 µg/ml doxycycline for 4 days to knock down endogenous BRD4-S and concurrently induce the expression of ectopic 3xFLAG-BRD4-S (1-719) WT or the YF-mutated protein. The cells were furthered incubated in serum-free DMEM for 24 hours before collected for cell viability and transwell migration assays as previously described [23]. Statistical significance was analyzed using GraphPad PRISM. Blots of samples from MDA-MB-231 transfected cells were probed using the following antibodies: 1:1000 anti-BRD4 N-terminus [23], 1:2000 M2 anti-FLAG (Sigma, F3165), 1:10000 anti-β-actin (Sigma, A1978).

## Results

### The tandem bromodomains of BRD4 associate more stably with poly-acetylated H4 tails than the individual bromodomains

Fully defined histone peptide and ‘designer’ nucleosome pull-down assays have been successfully used by our laboratory and others to interrogate the histone binding-preferences of chromatin readers [42–44]. A commonly used control protein for these assays is a GST fusion to the BD1 of BRD4, which robustly and preferentially binds poly-acetylated H4 tails [38]. Because GST naturally dimerizes and could artificially enhance apparent binding affinity, we sought to verify that BD1 alone was sufficient for poly-acetylated histone tail binding in our pull-down assays. After removing the GST tag by 3C protease cleavage and size exclusion chromatography (Fig. S1), we performed peptide pull-down assays using GST-BD1 and BD1 alone against unmodified or poly-acetylated H4 peptides. Since BD1 lacking a GST tag has been reported to robustly and preferentially bind to H4 poly-acetylated peptides and nucleosomes in other assay formats [25–27], we were surprised to find that untagged BD1 did not show detectable enrichment on either peptide in our pull-down assays, whereas GST-BD1 bound robustly to poly-acetylated and, to a lesser extent, unmodified H4 peptides (Fig. 1B). We therefore expanded this analysis to compare GST-BD1 and BD1 with BD2 and the tandem BD1-2 module. Neither untagged BD1 nor BD2 enriched on H4 peptides, whereas GST-BD1 and the untagged tandem BD1-2 module showed robust enrichment on poly-acetylated H4 peptides (Fig. 1B), suggesting that stable retention under stringent wash conditions is strongly enhanced by multivalency.

To rule out potential non-specific artifacts, we evaluated the folding of each purified BD1, BD2, and BD1-2 protein using differential scanning fluorimetry (DSF), where each protein displayed similar stabilities (Fig. S1). Further, we performed solution-based fluorescence polarization assays showing that BD1 and BD2 individually bind poly-acetylated H4 peptides, albeit weaker than the BD1-2 in tandem (Fig. 1C). This is as predicted of these bromodomains based on other solution based peptide binding experiments [25,26]. BD1-2 displayed a K_D_ of 13.79 ± 4.05 μM, compared to 30.18 ± 6.00 μM for BD1 and >200 μM for BD2 (Table 1). None of the constructs bound appreciably to the unacetylated peptide control (Fig. 1D). Although these precise affinity values differed from prior reports [25,26], the trends for BD1 and BD2 are largely consistent, with variations likely due to differences in the peptide modifications (*e.g.* 5-FAM) and acetylation status. Collectively, these results indicate that tandem BD1-2 combinatorially engages H4 poly-acetylated histone tails and is more stably retained under stringent biochemical conditions.

**Table 1.** Equilibrium binding values observed for BRD4 fluorescence polarization (FP) and biolayer interferometry (BLI) experiments (see Methods). H4 poly-acetylated peptide (H4_[1–23]_K5acK8acK12acK16acK20ac) and nucleosome ([H4K5acK8acK12acK16ac]_2_) were used in each FP and BLI, respectively. N/A for not applicable binding values and ND for curves that could not be accurately determined through fitting.

| Assay | BRD4 | Target | [BRD4] ($\mu\text{M}$) | $K_D$ ($\mu\text{M}$) | $K_D$ S.D. ($\mu\text{M}$) | $K_{D2}$ ($\mu\text{M}$) | $K_{D2}$ S.D. ($\mu\text{M}$) |
| --- | --- | --- | --- | --- | --- | --- | --- |
| Fluorescence Polarization | BD1-2 | **Peptide:** | N/A | 13.79 | 4.05 | N/A | N/A |
| | BD1 | $\text{H4}_{[1-23]}$ | N/A | 30.18 | 6.00 | N/A | N/A |
|  | BD2 | Kpoly-ac | N/A | 208.2 | 149.6 | N/A | N/A |
| Biolayer Interferometry |  | **Nucleosome:** | 12.5 | 1.89 | 0.06 | 23.9 | 1.4 |
| | BD1-2 | $[\text{H4Kpoly-ac}]_2$ | 6.25 | 0.48 | 0.03 | 1.58 | 0.05 |
|  |  |  | 3.13 | 0.90 | 0.02 | 6.70 | 0.24 |
|  |  |  | **Average:** | 1.09 | 0.04 | 10.74 | 0.84 |
|  | BD1-2 | Unmodified | ND | ND | ND | ND | ND |
| | BD1 | $[\text{H4Kpoly-ac}]_2$ | ND | ND | ND | ND | ND |
|  | BD1 | Unmodified | ND | ND | ND | ND | ND |

### The tandem bromodomains of BRD4 associate more stably with poly-acetylated H4 nucleosomes than the individual bromodomains

Previous high-throughput screens using DNA-barcoded nucleosome libraries demonstrated enhanced binding of the BRD4 tandem BD1-2 module over individual bromodomains [27]. To confirm and quantify this observation, we used biolayer interferometry (BLI), a quantitative kinetic binding method recently applied for studying histone reader-nucleosome interactions [45]. We observed robust, concentration-dependent binding of tandem BD1-2 on nucleosomes with both H4 tails poly-acetylated (*i.e.*, ([H4K5acK8acK12acK16ac]_2_), Fig. 2A), but not on unacetylated nucleosomes (Fig. 2B). Kinetic modeling (see methods and Fig. S2) yielded two distinct binding events with an average K_D_ of 1.09 ± 0.04 μM and 10.74 ± 0.84 μM, respectively (Table 1). The presence of multiple acetylated sites on the nucleosome could account for this biphasic binding behavior, with initial engagement followed by additional binding events within the same nucleosome (or possibly between nucleosomes) that enhance overall affinity and retention. Notably, BD1 alone showed significantly weaker or negligible nucleosome binding to either acetylated or unacetylated nucleosomes even at high protein concentrations (Fig. 2C and D). Together, these results indicate that BD1 and BD2 operating in tandem substantially increases BRD4’s capacity for efficient nucleosomal recognition, supporting the notion of combinatorial histone readout as crucial for BRD4 function.

**Figure 2.**
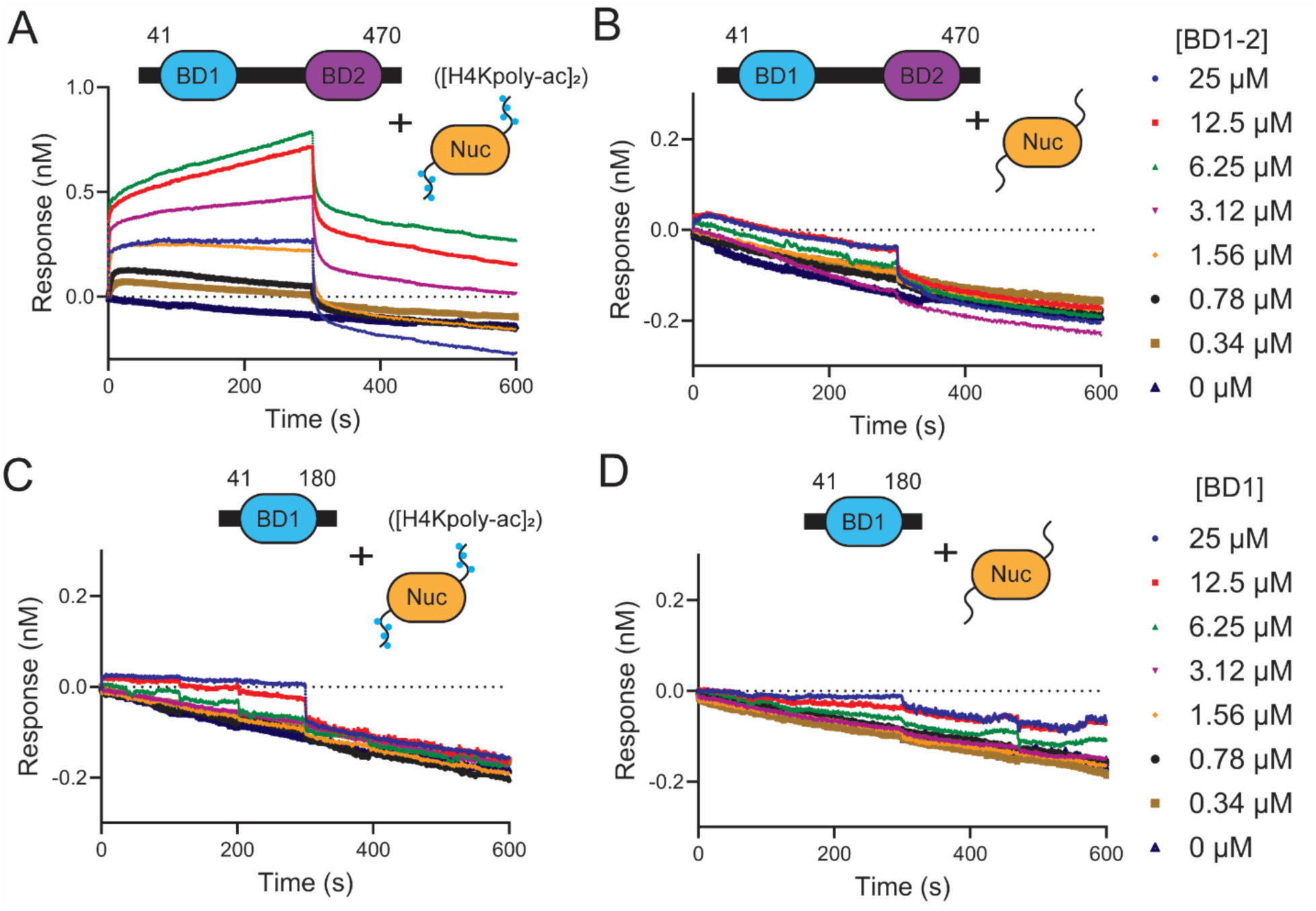
The BRD4 tandem bromodomains stably associate with H4 poly-acetylated nucleosomes in biolayer interferometry. (**A-D**) Biolayer interferometry binding assays of (**A**) BD1-2 with H4 poly-acetylated nucleosomes ([H4K5acK8acK12acK16ac]_2_), (**B**) BD1-2 with unacetylated nucleosomes, (**C**) BD1 alone with H4 poly-acetylated nucleosomes, and (**D**) BD1 alone with unacetylated nucleosomes. Protein concentrations are indicated at the right of each plot. Note the difference in y-axis scale between (**A**) 1 to -0.5 nm and (**B-D**) 0.3 to -0.3 nm. Octet fitting of BD1-2 binding curves to H4 poly-acetylated nucleosomes (12.5, 6.25, and 3.12 μM) using a 2:1 heterogeneous ligand model produced R^2^ values >0.98 (Fig. S2). Experiments were performed in triplicate.

As BRD4-nucleosome binding behavior can vary with construct and assay format [27,34], we also tested the binding of GST-tagged BRD4 BD1-2 to a panel of designer nucleosomes using a high-throughput, bead-based Luminex platform (also known as Captify^TM^-Luminex) [46,47]. Consistent with prior reports showing BRD4 can associate with nucleosomes in an acetylation-independent manner at lower ionic strength [34], GST-BD1-2 displayed strong, largely nonselective binding to nucleosomes under low-salt conditions (Fig. S3A). In sharp contrast, increasing NaCl concentration to 150 mM revealed preferential binding of the GST-BD1-2 to H4 poly-acetylated nucleosomes ([H4K5acK8acK12acK16ac]_2_) relative to the others tested; with decreasing binding across H3 poly-acetylated ([H3K4acK9acK14acK18ac]_2_), tailless, H4K12 acetylated ([H4K12ac]_2_), H4 N-terminal tail truncated ([H4ΔN15]_2_), and unmodified nucleosomes (Fig. S3B). At 225 mM NaCl, robust binding was observed only for H4 poly-acetylated nucleosomes with weak residual binding to H3 polyacetylated nucleosomes, and barely detectable signal for the others in the panel (Fig. S3C). Since altering the salt concentration could affect nucleosomal stability in these assays, we used a pan-histone antibody to monitor bead-bound nucleosome levels across each condition. Increasing salt concentration does lead to a modest reduction in detectable histone levels in this stringent wash-based method, but the drop was relatively consistent for each nucleosome and clearly not reflective of the stark change in GST-BD1-2 specificity at higher salt concentration (Fig. S3D). Additionally, it is difficult to discern whether a drop in histone signal at higher salt concentrations is due to nucleosomal collapse and other factors such as potentially altering the antibody’s affinity for histones. Together, these results indicate that the tandem BRD4 bromodomains can non-specificially engage nucleosomes at low ionic strength, but become highly selective for H4 poly-acetylated nucleosomes at near-physiological salt concentrations [48,49].

### The tandem bromodomains of BRD4 associate more stably with chromatin in cells than the individual bromodomains

Given the significant distinction between the tandem and individual bromodomains in our histone peptide and nucleosome binding experiments *in vitro*, we next investigated the functional consequences in cells. We generated and stably integrated 3xFLAG-tagged, Tet-inducible mammalian expression constructs for BD1, BD2, and BD1-2 in HEK293T cells. We observed modest chromatin association of BD1 alone, while BD2 alone had negligible chromatin association (Fig. 3A and B). In contrast, the tandem BD1-2 module had robust chromatin association that greatly exceeded the individual domains (Fig. 3A and B).

**Figure 3.**
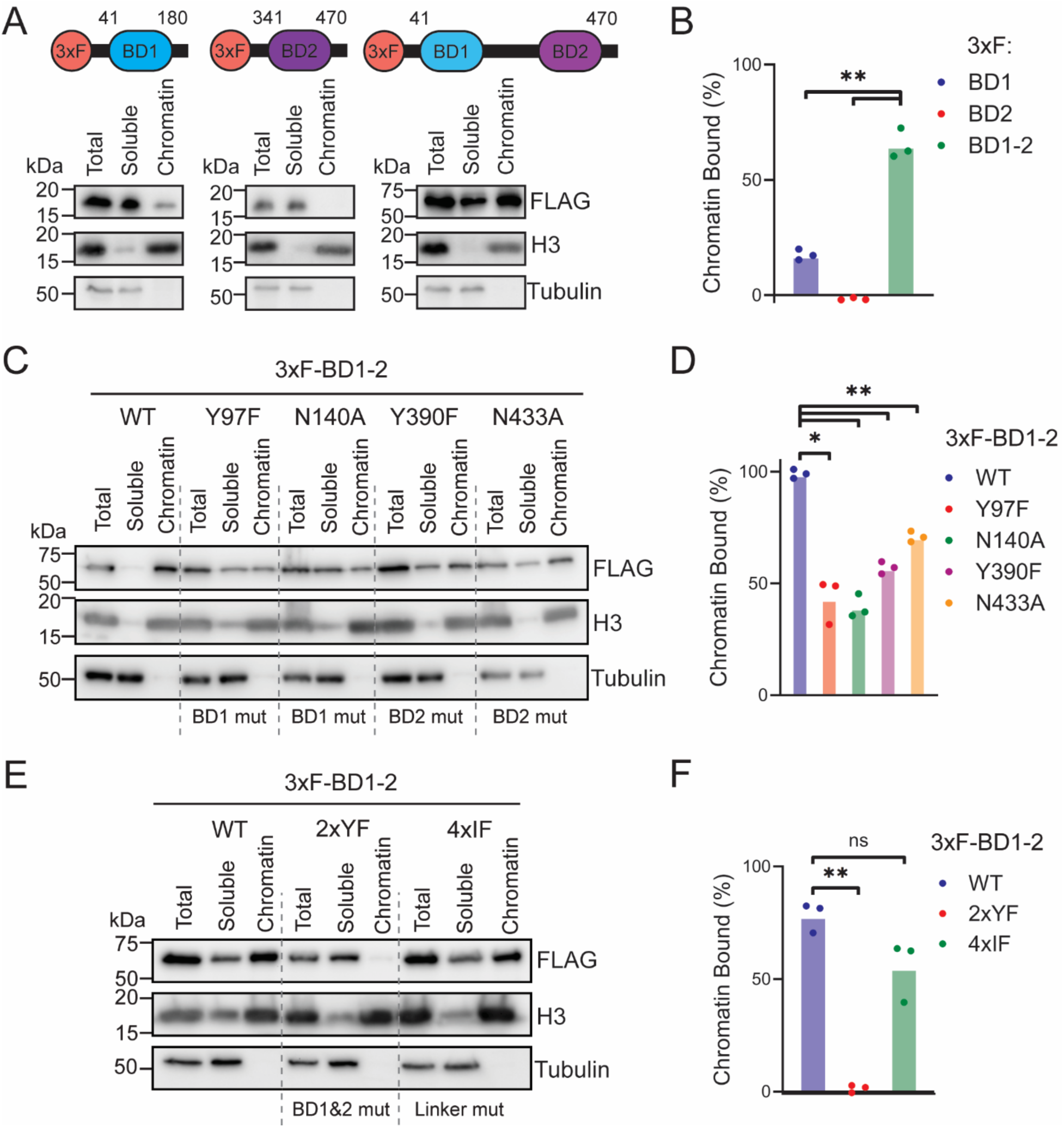
The BRD4 tandem bromodomains robustly associate with chromatin in cells. (**A**) Chromatin association assays of indicated 3xFLAG-tagged BRD4 constructs expressed in doxycycline-induced HEK293T cells. Western blots were probed with antibodies against FLAG (BRD4), pan-histone H3 (chromatin fraction control), and α-tubulin (soluble fraction control). (**B**) Quantification of mean chromatin-to-soluble protein ratios by densitometry from three biological replicates. Statistical significance was calculated by two-tailed t-tests (*p<0.05, **p<0.005). (**C-D**) Chromatin association assays and densitometry quantifications using indicated 3xFLAG-tagged BRD4 BD1-2 constructs with mutations in individual bromodomain acetyl-binding pockets averaged from three biological replicates. (**E-F**) Chromatin association assays and densitometry quantifications testing dual BD1-2 mutants: 2xYF (Y97F+Y390F, disrupting acetyl-binding pockets) and 4xIF (Q78A, F79A, Y372A, K445A; residues implicated in interdomain interface) averaged from three biological replicates.

To dissect the contributions of individual bromodomains within the tandem module, we introduced point mutations in the acetyl-binding pockets of BD1 (Y97F, N140A) or BD2 (Y390F, N433A) that disrupt acetylated histone recognition [26]. These mutated forms had diminished chromatin association compared to the wild-type BD1-2 (Fig. 3C and D). The BD1 mutations had a slightly stronger impact on chromatin association than the BD2 mutations (Fig. 3C and D). Notably, despite BD2 alone exhibiting negligible chromatin association (Fig. 3A and B), BD2 pocket mutations in the BD1-2 construct caused defects comparable to BD1 pocket mutations (Fig. 3C and D). This suggests that while BD1 contributes substantially to chromatin association, acetyl-reading by the BD2 and DNA binding by the linker and BD2 become more important in the context of the tandem BD1-2 module in cells.

To further clarify the role of both BDs, we generated constructs with dual mutations (Y97F + Y390F), in the BD1 and BD2 domains (2xYF). Comparison of 2xYF with wild-type BD1-2 in cells demonstrated a drastic loss of chromatin association when both domains were mutated (Fig. 3E), indicating that combinatorial acetyl recognition by both domains is required for stable chromatin association in this cellular context. Additionally, we explored the potential role of BRD4 dimerization on chromatin binding given previous studies that identified a putative binding interface within the bromodomains from crystallographic experiments [25]. Mutation of these interface residues (Q78A, F79A, Y372A, K445A; 4xIF) only marginally affected chromatin association, suggesting that direct interdomain interactions are less critical than dual-domain acetyl recognition (Fig. 3E and F). Finally, given that linker region mutations can be disease associated and compromise histone reading (*e.g.*, PHIP R1310C [29]) we tested BRD4 linker mutation Q209H, a frequently identified allele in the COSMIC database of cancer mutations. However, this mutation did not cause a significant defect in chromatin association in our assays (Fig. S4), reinforcing the concept that the primary determinant of efficient chromatin targeting in this system is the dual-bromodomain acetyl recognition. Together, these results indicate that the BRD4 bromodomains being in tandem facilitates a cooperative mechanism that promotes chromatin association.

### Both bromodomains are required for efficient BRD4-S chromatin association

Given that bromodomain inhibition is widely used to disrupt BRD4 function, it was important to determine whether its bromodomains are similarly required for chromatin association in a more biologically relevant context. We therefore tested an engineered BRD4-S (1-719) construct bearing an N-terminal 3xFLAG tag and lacking the three C-terminal residues (GPA) unique to endogenous BRD4-S (Fig. 4A). We focused on BRD4-S because it has been implicated in promoting oncogenic phenotypes in a triple negative breast cancer model in contrast to BRD4-L [23]. Wild-type BRD4-S (1-719) was able to robustly bind to chromatin as expected (Fig. 4B and D). Unexpectedly, single mutations in either bromodomain did not appear to disrupt BRD4-S (1-719) chromatin association, unlike what we observed with the BD1-2 construct (Fig. 3E and F). On the other hand, mutation of both domains yielded ∼60% reduction in chromatin-associated BRD4-S (1-719) (Fig. 4B and D), a contrast to the ∼90% reduction observed for the 2xYF BD1-2 construct (Fig. 3E and F). These findings suggested that the tandem BD1-2 module is the major driver of BRD4-S chromatin association, but additional regions also contribute. To examine this further, we created N-terminal and C-terminal truncations (respectively BRD4-S (1-440) 2xYF and BRD4-S (75-719) 2xYF, Fig. 4A), observing the latter to display a significant drop in chromatin association (Fig. 4C and D). Together, these experiments indicate that the C-terminal portion of BRD4-S cooperates with the tandem bromodomains to promote chromatin engagement.

**Figure 4.**
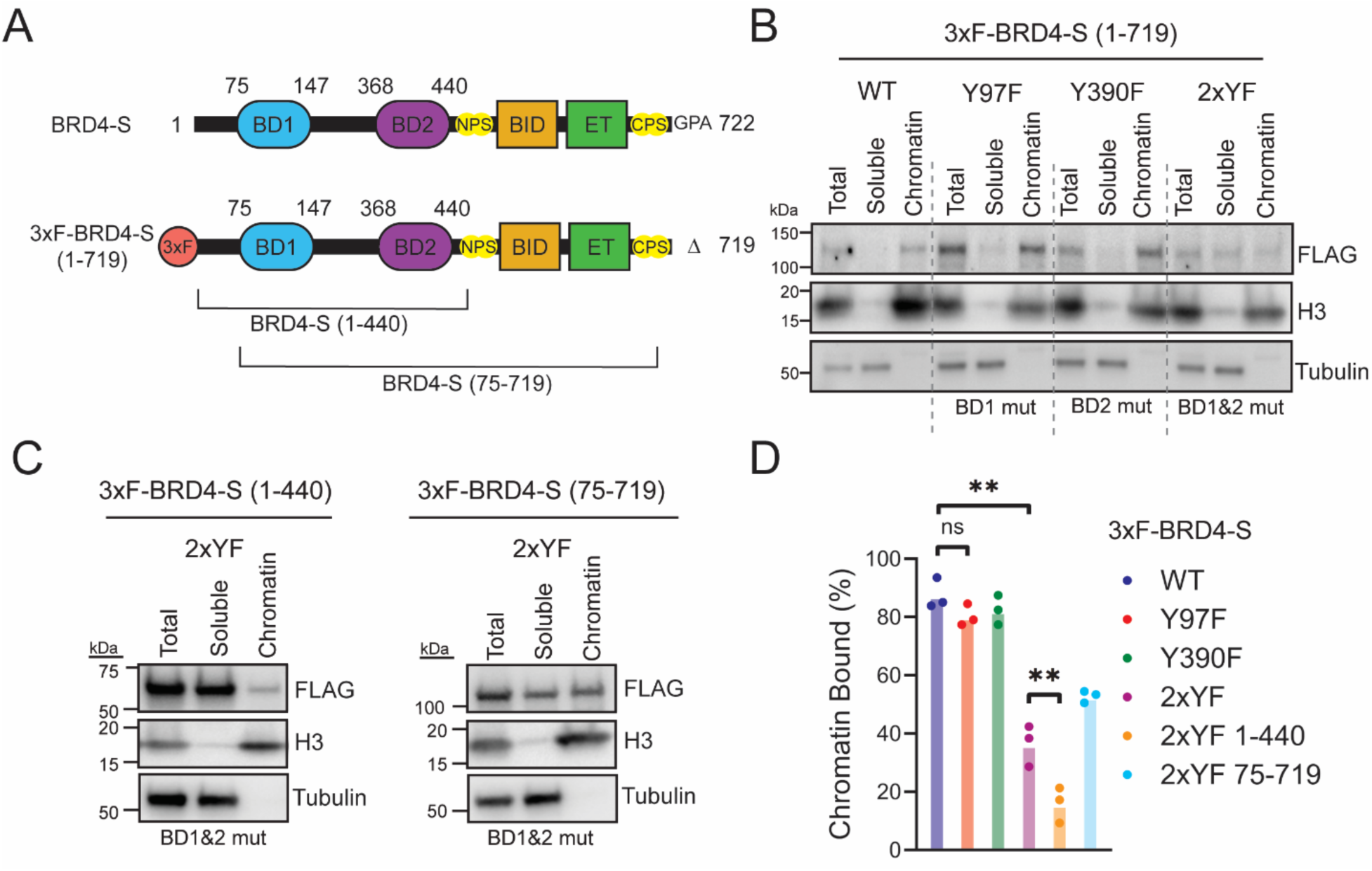
Both bromodomains and the C-terminal region of engineered BRD4-S construct are required for maximal chromatin association in cells. (**A**) Schematic of endogenous BRD4-S and engineered 3xFLAG-BRD4-S (lacking the unique C-terminal GPA sequence) along with truncation mapping below. (**B**) Chromatin association assays of wild-type, single pocket mutant, or double pocket mutant (2xYF) BRD4-S expressing cells. (**C**) Chromatin association assays of BRD4-S 2xYF truncations (1-440 and 75-719). (**D**) Quantification of chromatin-to-soluble band densities from chromatin association assays in (**B-C**). All experiments were run in biological triplicate. Statistical significance was calculated by two-tailed t-tests (*p<0.05, **p<0.005).

### An AlphaFold model suggests a plausible model of BRD4-S engagement with an H4K12-acetylated nucleosome

As no experimental structure of BRD4 with both bromodomains bound to a nucleosome has yet been resolved, we used AlphaFold to generate a hypothesis-generating structural model of BRD4S in complex with an H4 acetylated nucleosome. We initially attempted to model BRD4-S bound to a poly-acetylated H4 nucleosome, but AlphaFold was unable to produce an interpretable model for this more complex substrate. We therefore simplified the input and modeled BRD4-S bound to a nucleosome containing H4K12 acetylation on each H4 tail, guided in part by a recent cryo-EM structure showing BRD4 BD1 bound to a H4K12, H4K16, and H3K18 acetylated nucleosome ([H3K18ac]_2_·[H4K12acK16ac]_2_) [35]. In the top-ranked model, BD1 and BD2 independently engage the two H4K12ac tails while remaining linked by a flexible intervening region that tracks along the nucleosomal DNA (Fig. 5A and B). Although this model should be interpreted cautiously, it shows that intranucleosomal co-engagement by linked BRD4 bromodomains is physically plausible and provides a possible structural rationale for the enhanced nucleosome binding we observed for the tandem BD1-2 module. The model also suggests that residues outside the acetyl-lysine binding pockets, including basic residues near BD2, could contribute to nucleosome engagement through contacts with DNA. At the same time, the model does not explain our finding that the C-terminal region of BRD4-S promotes chromatin association in cells indicating that additional conformations, chromatin binding modes, or partner-mediated interactions are likely involved. Thus, we view this model primarily as a framework for future testing rather than as definitive evidence for a single BRD4S–nucleosome binding mode. We also emphasize that this model does not preclude the potential for BRD4 bromodomains to engage distinct nucleosomes, as has been recently suggested [50].

**Figure 5.**
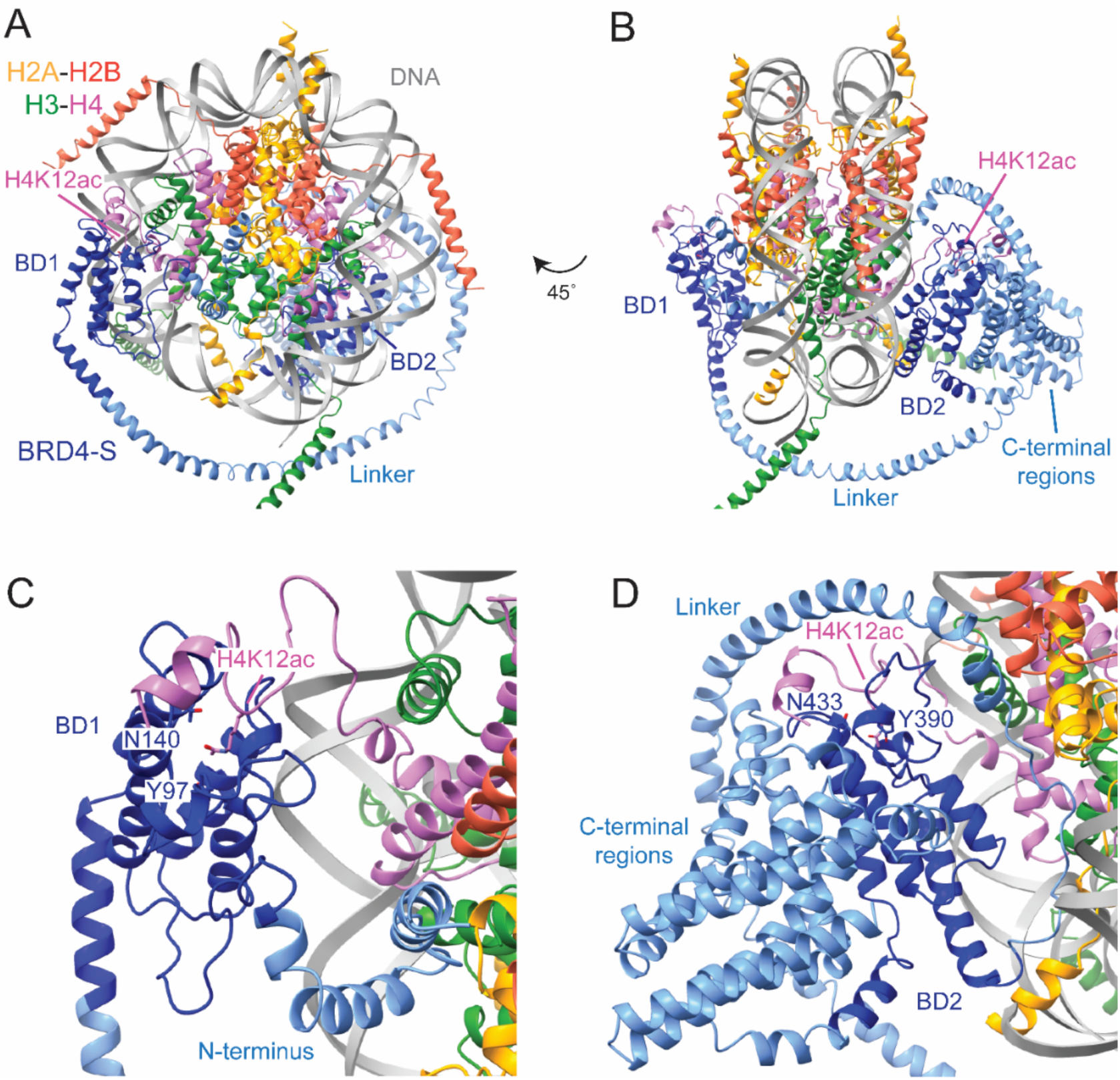
AlphaFold model suggesting a plausible mode of BRD4-S engagement with an H4 K12 acetylated nucleosome ([H4K12ac]_2_). (**A**) Top-ranked AlphaFold model prediction of BRD4-S (blue) in complex with an H4K12ac nucleosome (H2A, golden; H2B, tomato; H3, green; H4, violet, Widom 601 DNA, grey). BD1 and BD2 are shaded in darker blue for clarity. (**B**) Rotated (45°) view of the modeled BRD4-S-H4K12ac nucleosome complex. (**C**) Zoom-in of the BD1 acetyl-lysine binding pocket highlighting residues Y97 and N140 positioned near H4K12ac. (**D**) Zoom-in of the BD2 acetyl-lysine binding pocket highlighting residues Y390 and N433 positioned near H4K12ac. This view also shows the relevant placement of the C-terminal BRD4-S region including the BID, ET, and phosphoregulatory sites, in the model.

In the AlphaFold BRD4-S-bound nucleosome model, both bromodomains engage each H4K12ac through their aromatic cages with the BD1 binding to the first K12ac with residues Y97 and N140 (Fig. 5C) while BD2 engages the second K12ac on the other side of the nucleosome through residues Y390 and N433 (Fig. 5D). The model places basic residues of BD2, including H341 and K355, in proximity to nucleosomal DNA (Fig. S5), consistent with recent studies implicating DNA contacts as an important component of BRD4-nucleosome engagement [34,35]. However, the C-terminal regions of BRD4-S are not positioned near the nucleosome in the model, despite our truncation data indicating that this region contributes to chromatin association (Fig. 4C and D). This mismatch underscores the limitations of the current model and suggests that BRD4-S may adopt alternative conformations or that the C-terminus is engaging additional chromatin associated factors not captured here. Together, this model provides a possible view of how BRD4-S can engage H4 acetylated nucleosomes that may help guide future exploration of chromatin engagement and potential druggable interaction surfaces.

### Mutation of the bromodomain pockets in BRD4-S attenuate breast cancer growth and migration phenotypes

Given the broad interest in therapeutic targeting BRD4 bromodomains in cancer [13,14,22,51], we tested the impact of bromodomain pocket mutations in a relevant cancer model system using the TNBC cell line MDA-MB-231. Depletion of BRD4-S in this cell line reduces cell viability and oncogenic properties such as the capacity to migrate through semi-permeable surfaces [23]. We introduced 3xFLAG-tagged BRD4-S (1-719) wild-type and BD pocket mutants into a version of this cell line containing doxycycline-inducible shRNAs targeting endogenous BRD4-S 3’ UTR sequences [23]. Thus, we were able to simultaneously deplete endogenous BRD4-S while complementing with wild-type or mutant 3xFLAG BRD4-S (1-719) through addition of doxycycline. Complementation with wild-type BRD4-S (1-719) restored much of the viability defect from endogenous BRD4-S knockdown (Fig. 6A), indicating that the slightly truncated BRD4-S (1-719) construct functionally mimics endogenous BRD4-S. Interestingly, the BD2 pocket mutant complemented viability, whereas the BD1 and 2xYF mutants did not (Fig. 6A), implicating BD1 as being particularly important for this function of BRD4-S.

**Figure 6.**
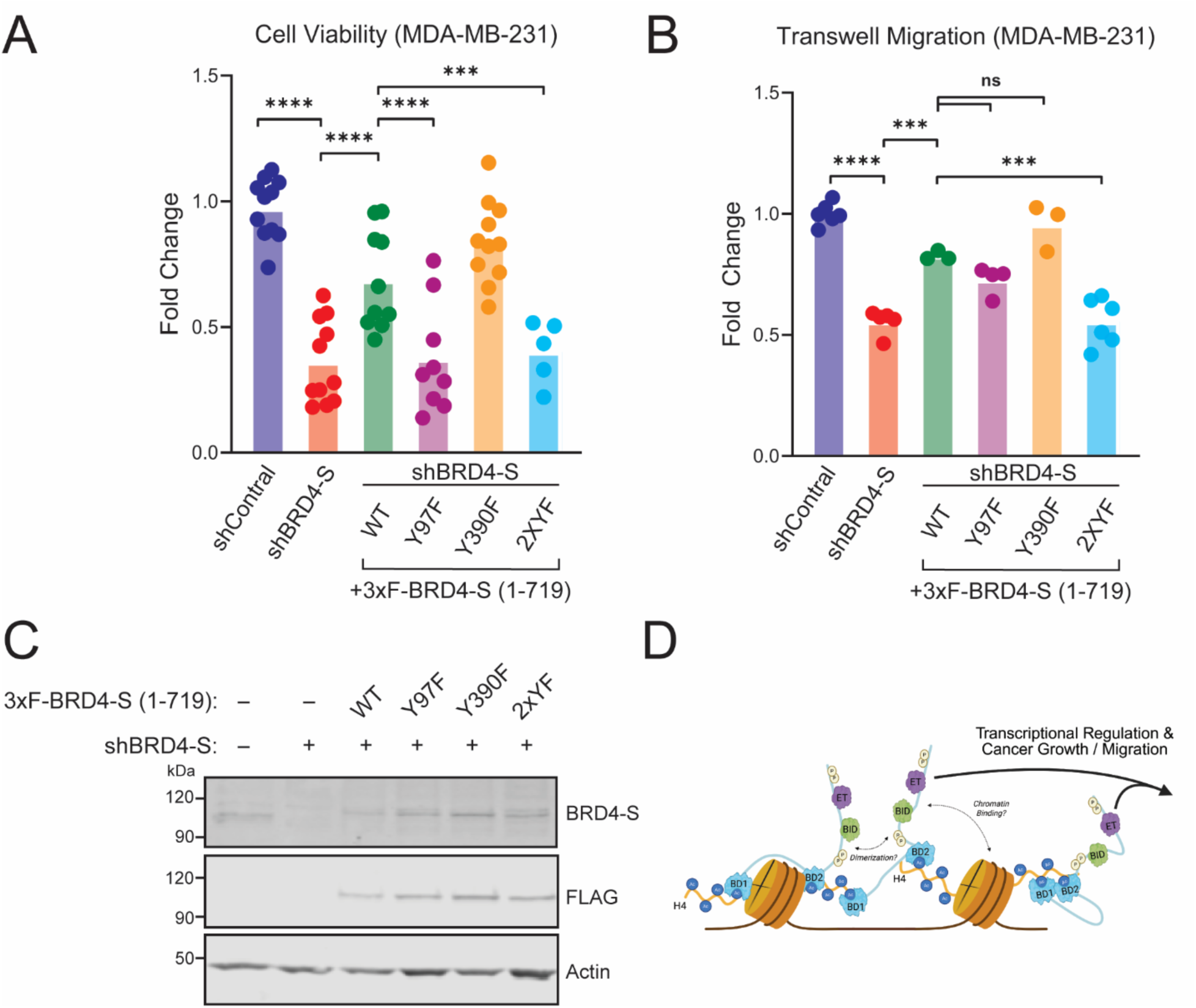
Bromodomain binding pocket mutations in the BRD4-S disrupt breast cancer growth and migration phenotypes. (**A**) Cell growth assay following BRD4-S depletion and complementation with the indicated 3xFLAG-BRD4-S constructs (mean ± SEM; n≥5 independent experiments). Statistical analysis as described in Methods. (**B**) Transwell migration assay following BRD4-S depletion and complementation with the indicated 3xFLAG-BRD4-S constructs (mean ± SEM; n ≥ 3 independent experiments). Statistical significance was calculated by two-tailed t-tests (***p<0.0005, ****p<0.0001). (**C**) Western blots of endogenous BRD4-S depletion and rescue by 3xFLAG-tagged BRD4-S (1-719) constructs in doxycycline-treated MDA-MB-231 cells. Blots were probed with an antibody raised for the N-terminus of BRD4 (BRD4-S; for endogenous ∼100 kDa), M2 FLAG (FLAG), and β-actin (Actin) (**D**) Graphical illustration of putative model for BRD4-S chromatin engagement and regulation of cancer cell growth and migration (BioRender, https://BioRender.com/7dfa8yg). This model shows multiple possible binding modes of BRD4-S on poly-acetylated nucleosomes that would be difficult to distinguish or determine in biochemical and structural studies, yet could all contribute to recruitment of regulatory complexes and transcriptional control of nearby genes including those for cancer growth and migration.

In transwell migration assays, wild-type BRD4-S (1-719) largely complemented BRD4-S-dependent migration across semi-permeable surfaces (Fig. 6B). While individual pocket mutants showed partial effects, only the 2xYF mutant was fully incapable of complementing the BRD4-S-dependent migration phenotype (Fig. 6B), indicating that both bromodomains in tandem contribute to this phenotype. Together, these experiments suggest that both bromodomains are critical for the oncogenic functions of BRD4-S in this TNBC cell model, even though the BD1 appears to be more important for viability under these conditions. Although some variability in BRD4-S (1-719) expression was observed across experiments, expression of the constructs was generally comparable and does not explain the distinct complementation patterns observed for viability and transwell migration (Fig. 6A-C).

Altogether, our results support a model in which BRD4-S engages hyperacetylated H4-marked chromatin in a multivalent fashion via its tandem bromodomains and C-terminus to regulate transcriptional and cellular functions, including breast cancer cell growth and migration (Fig. 6D). It is important to note that these results pertain specifically to BRD4-S function in a 2D breast cancer cell model as the two bromodomains could be functioning differently in other cancer types and molecular contexts such as during DNA damage response [52].

## Discussion

Through a series of stringent wash-based peptide and nucleosome binding assays performed in parallel with cell-based chromatin association assays, we found that the bromodomains of BRD4 cooperatively engage chromatin in ways that support its cellular functions. Recent work has highlighted that BRD4-nucleosome interactions can be strongly influenced by protein-DNA contacts and, in some settings, show only modest dependency on acetyl-histone binding [34,35] including in our own low salt Luminex-based nucleosomal binding experiment (Fig. S3). In our assays, however, the BD2 contributes substantially to H4 acetylated histone peptide binding, acetylated H4 nucleosome engagement, and cellular chromatin association when operating in tandem with the BD1 under physiological ionic conditions. We suspect that BD2 supports these functions through a combination of: (1) weaker acetyl-histone binding (Fig. 1C); (2) DNA contacts [34,35] that are likely very sensitive to ionic conditions; and (3) combinatorial readout with BD1 to remain more tightly associated with poly-acetylated H4 (Fig. 5A and B). These properties of BRD4 are all consistent with a multivalent mechanism in which DNA interactions provide a basal tether while tandem bromodomains enhance acetyl-dependent avidity and chromatin residence. A caveat is that the BRD4 bromodomains can also recognize non-histone targets such as RelA [53,54] and TWIST1 [55] as well as potentially other chromatin-associated proteins and/or noncoding RNAs in cells [56,57], which could influence BRD4’s overall capacity to associate with chromatin. Nevertheless, binding to acetylated transcription factors is also perturbed by inhibition of the BRD4 bromodomains with JQ1 [54,55], supporting the relevance of our results for understanding how bromodomain targeting impacts BRD4 chromatin association and downstream biology. Altogether, our data supports a model in which BRD4 employs a multidomain strategy for engaging chromatin and exerting its various biological functions [51,58].

In addition to elucidating the cooperative role of tandem bromodomains, we uncovered a chromatin-associating function within the BRD4 sequence between BD2 and the end of the BRD4-S isoform (Fig. 4B-D). Several functional regions reside in this segment, including N- and C-terminal clusters of phosphorylation sites (NPS: aa 485-504; CPS: aa 699-717), a basic residue-enriched interaction domain (BID: aa 503-548), and an extraterminal domain that mediates protein-protein interactions (ET: aa 601-679; Fig. 1A). One attractive explanation for why this region enhanced chromatin association in our assays is NPS phosphorylation-dependent intermolecular homodimerization with endogenous and/or exogenous BRD4-S proteins with intact bromodomains [17,59]. Normally, the NPS interacts with the adjacent BD2 reducing its capacity for recognizing acetylated histones [17], but phosphorylation of NPS releases it from BD2 and allows it to interact either with the adjacent BID or a BID from another protein such as other BRD4 molecules [59]. BRD4 homodimerization could increase local avidity by enabling multiple BRD4 molecules, each containing two bromodomains, to cooperatively assemble at sites of histone hyperacetylation such as enhancers [60]. Given that BID-dependent dimerization is also promoted by phosphorylation of the neighboring NPS region [17,59], BRD4 hubs could be formed and dissolved quickly through dynamic reversible phosphorylation at genomic loci enriched in H4 poly-acetylation and then deacetylated to turn off transcription regulation (Fig. 6D). Notably, only a tandem BRD4 bromodomain inhibitor appeared to significantly disrupt BRD4 dimers compared to mono-bromodomain inhibition [59], supporting a view in which bromodomain engagement and dimerization-mediated avidity cooperate to stabilize chromatin association.

Additional layers of chromatin targeting may also arise through regulated protein interactions in the C-terminal region of BRD4-S, including p53 binding when the NPS is dephosphorylated which is essential for BRD4’s recruitment and activation of p53-targeted gene transcription [17]. Likewise, the BRD4 ET can associate with several transcriptional regulators, including the short isoform of NSD3 (NSD3S), JMJD6, CHD4, GLTSCR1, and ATAD5 [61,62], which could further enhance BRD4 chromatin association and functional specificity. Importantly, inhibition of the BRD4 bromodomains, NSD3S, or CHD8 disrupts BRD4-NSD3S-CHD8 complexes at super-enhancers in acute myeloid leukemia cells and reduces their proliferation [62], highlighting the functional importance of bromodomain-mediated chromatin engagement within oncogenic transcriptional networks. Additionally, BRD4 transcriptionally cooperates at enhancers of activated macrophage immune genes with GLTSCR1 and BRD9 containing noncanonical BAF remodeling complexes highlighting the complexity of BRD4’s roles [63]. Based on these connections, an important next step will be to define how the different BRD4 regions, including the lesser understood CPS and the full C-terminal sequence of BRD4-L, contribute to chromatin association through more focused mutation, truncation, and inhibitor-based experiments to determine whether their contributions reflect additional direct chromatin contacts, recruitment of partners, or both.

Our work exploring the histone and chromatin binding regions of BRD4 also motivated us to examine their roles in BRD4-S-promoting oncogenic phenotypes in breast cancer (Fig. 6). In MDA-MB-231 TNBC cells depleted of endogenous BRD4-S, we observed that the viability defect was most sensitive to complementation with BRD4-S constructs carrying a BD1 pocket mutation (Fig. 6A). In contrast, only the doubly mutated BD1/BD2 construct failed to restore migration across transwell inserts (Fig. 6B), a phenotype relevant to invasive behavior. The basis for this discrepancy remains unclear, but one possibility is that distinct gene programs have different BRD4 dosage or chromatin residence requirements (*e.g.*, growth-related transcriptional programs versus extracellular matrix and matrisome programs implicated in migration and metastasis) [23]. Defining these differences will require further genomic and mechanistic follow-up studies. More broadly, as BRD4 has been linked to many cancers and is of continued interest for BET inhibitor treatment strategies [21,51], our findings emphasize the need to better understand the biological consequences of disrupting specific BRD4 interactions including histone tails, nucleosomal DNA, and protein-protein interactions on chromatin [58]. This is especially true in the sense that the various BRD4 domains have been linked to unique functions in regulating BRD4 activity in differing cancer types [52]. Optimization from first generation BET inhibitors has produced family member-specific compounds [64,65], bromodomain-specific compounds [66], and even tandem-bromodomain targeting compounds [67]. Our results support the use of tandem inhibitors when the goal is maximal disruption of BRD4 chromatin engagement, but they also suggest that inhibition strategies may require nuanced approaches, as different cancer phenotypes (*e.g.*, viability and migration) may exhibit distinct sensitivities to disruption of one or both bromodomains [68]. This is especially important given that sustained BRD4 inhibition can lead to toxicity in mammals [69]. Thus, careful selection, dosage, and mechanistic matching of BRD4 BET inhibitors to disease context will likely be required for successful translation in human trials.

Finally, several emerging therapeutic alternatives may compliment BRD4 bromodomain inhibition, including recently identified phospho-NPS and ET targeting inhibitors [58,70–72]. These compounds respectively interfere with BRD4 homodimerization and key protein-protein interactions, potentially without perturbing BRD4’s ability to bind hyperacetylated histones and chromatin. These strategies could, in principle, reduce side effects by allowing BRD4-L to maintain parts of its chromatin-associated transcriptional functions while selectively blocking the oncogenic effects of BRD4-S, though this will require careful evaluation in relevant models. In the context of our findings, these strategies are particularly interesting because they target regions that appear to contribute to BRD4-S chromatin association and functional specificity beyond the bromodomains. Thus, it would be very interesting to explore how the BID and ET regions, and the regulatory phosphorylation sites flanking them, shape BRD4 chromatin engagement and biological function for informing future cancer treatment strategies.

## Supporting Information

Supplementary data is available at NAR online

## Acknowledgements

We thank Dr. Ashutosh Tripathy, Director of the UNC Macromolecular Interactions Facility, for assistance in performing the biolayer interferometry experiments. We thank Dr. Michael-Christopher Keogh, Chief Scientific Officer at EpiCypher, for providing constructive feedback on the manuscript.

## Author Contributions

N.T.B. and B.D.S. conceptualized the project and co-wrote the manuscript. N.T.B. and J.H. performed the *in vitro* binding experiments with support from I.K.P. and M.R.M. in the Captify^TM^ experiments. K.K. generated the histone peptides. N.T.B., J.H., and C.B. performed the chromatin association assays. S.-Y.W. and C.-M.C. conceptualized the breast cancer experiments and S.-Y.W. performed them, with both also involved in editing the manuscript.

## Data Availability

The authors confirm that the data supporting the findings of this study are available within the article and its supplementary information. Raw data files are available upon request and original Western blot images used in this study are presented in Fig. S6.

