## Supplemental data for "Tandem bromodomains of BRD4 cooperatively read poly-acetylated nucleosomes to enhance chromatin engagement and regulate breast cancer phenotypes"

**Supplemental Information – Burkholder et al.**

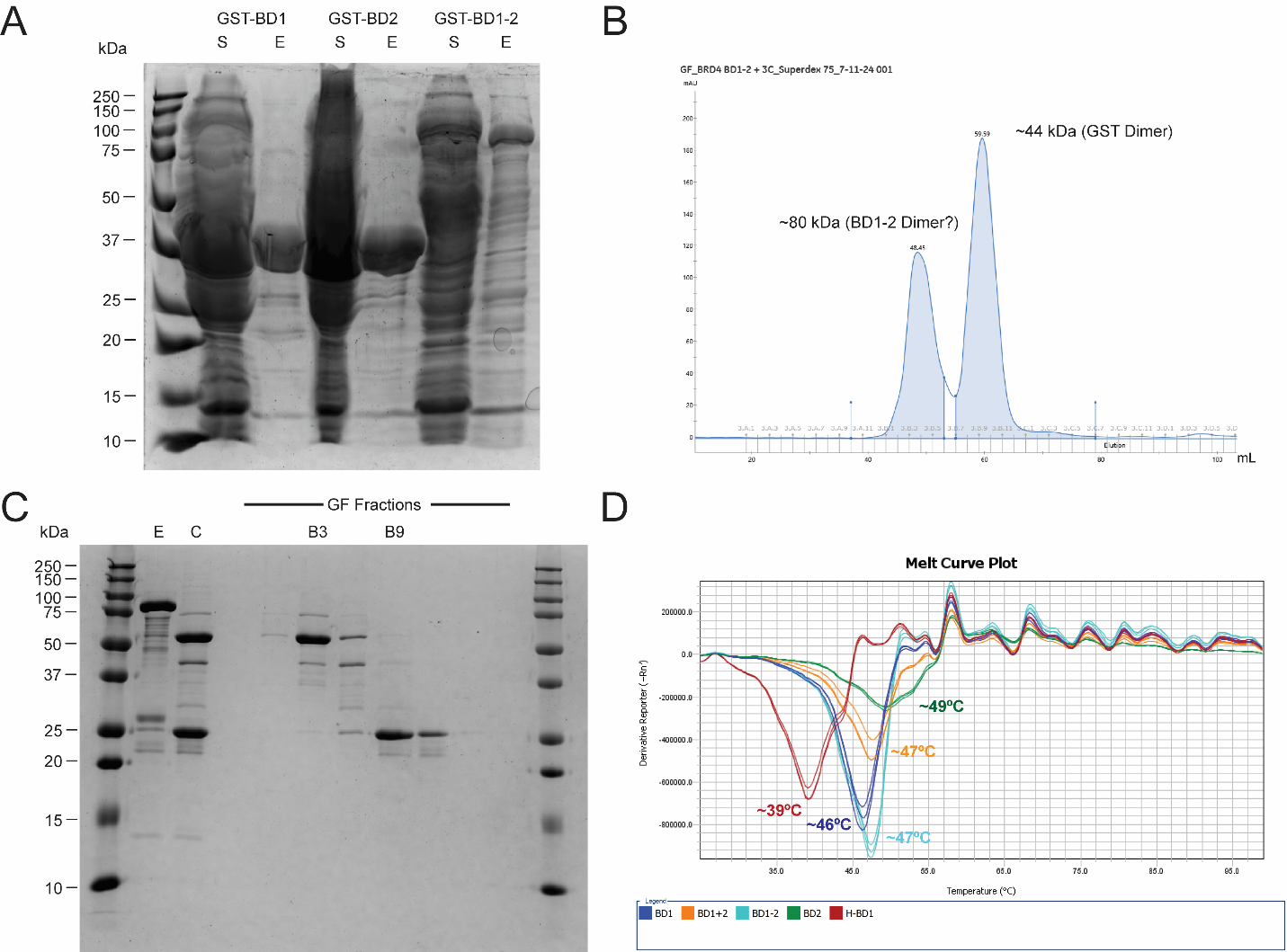

**Supplemental Figure S1.** Purification of BRD4 bromodomain constructs. (**A**) Soluble (S) and elution (E) samples from GST purifications analyzed by Coomassie-stained SDS-PAGE. Calculated molecular weights are as follows: GST-BD1-2, ~75 kDa; GST-BD1, ~43 kDa; GST-BD2, ~42 kDa. (**B**) Size exclusion chromatography of 3C-cleaved BRD4 BD1-2 (~49 kDa) over a Superdex 75 pg column (Cytiva). Calculated molecular weights for major peaks (based on a standard curve) indicated above. BD1-2 and GST appear to elute at an apparent size consistent with dimerization under these conditions. Similar chromatography was performed for the 3C-cleaved BD1 and BD2, which had similar purification quality as BD1-2 (data not shown). (**C**) Coomassie-stained gel of BD1-2 elution (E), 3C-cleaved (C), and gel filtration (GF) fractions from Fig. S2B. Fraction B3 was collected and concentrated as final BD1-2 purified protein. A similar approach was performed for the BD1 and BD2 samples. (**D**) Differential scanning fluorimetry of purified BD1-2 (teal), BD1 (blue), and BD2 (green) proteins. Approximate melting temperatures (Tm) averaged from triplicates are indicated. 6xHis-tagged BD1 (red) and pooled unlinked BD1 + BD2 (orange) were also purified and tested by peptide pull-down assays but exhibited no specific binding to poly-acetylated peptides (data not shown).

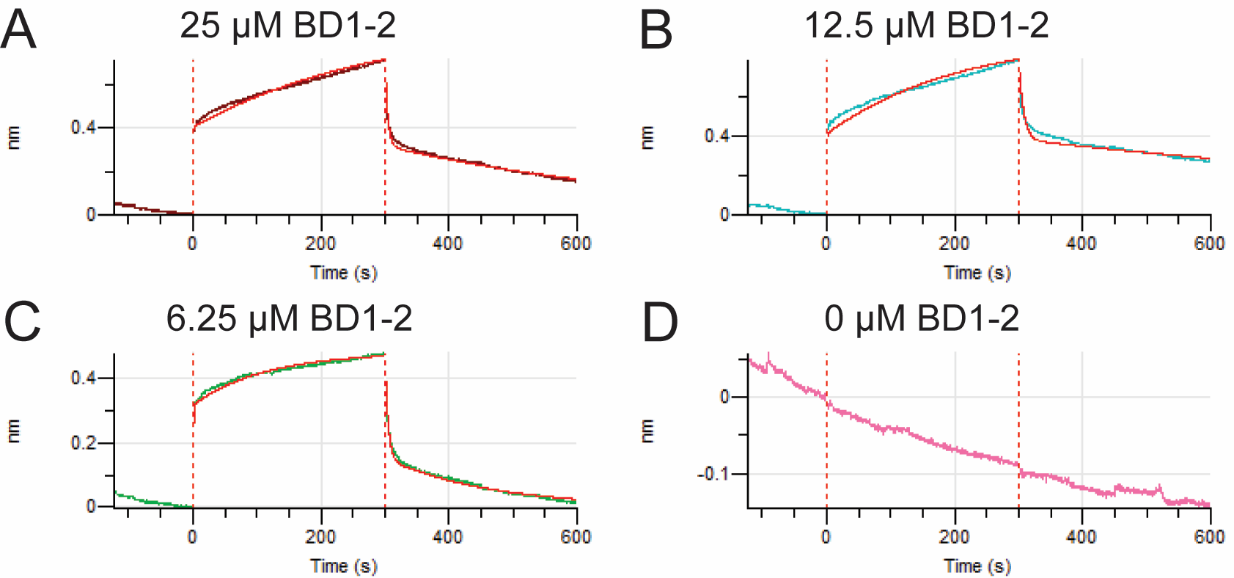

**Supplemental Figure S2.** BD1-2 binding to H4 poly-acetylated nucleosomes detected by biolayer interferometry. (**A-D**) Octet fitting curves (2:1 heterogenous ligand model, Sartorius) of biolayer interferometry data for BD1-2 binding to H4 poly-acetylated nucleosomes ([H4K5acK8acK12acK16ac]_2_) at concentrations of (**A**) 25 μM (R^2^ = 0.9953), (**B**) 12.5 μM (R^2^ = 0.9809), (**C**) 6.25 μM (R^2^ = 0.9982), and (**D**) control (0 μM BD1-2).

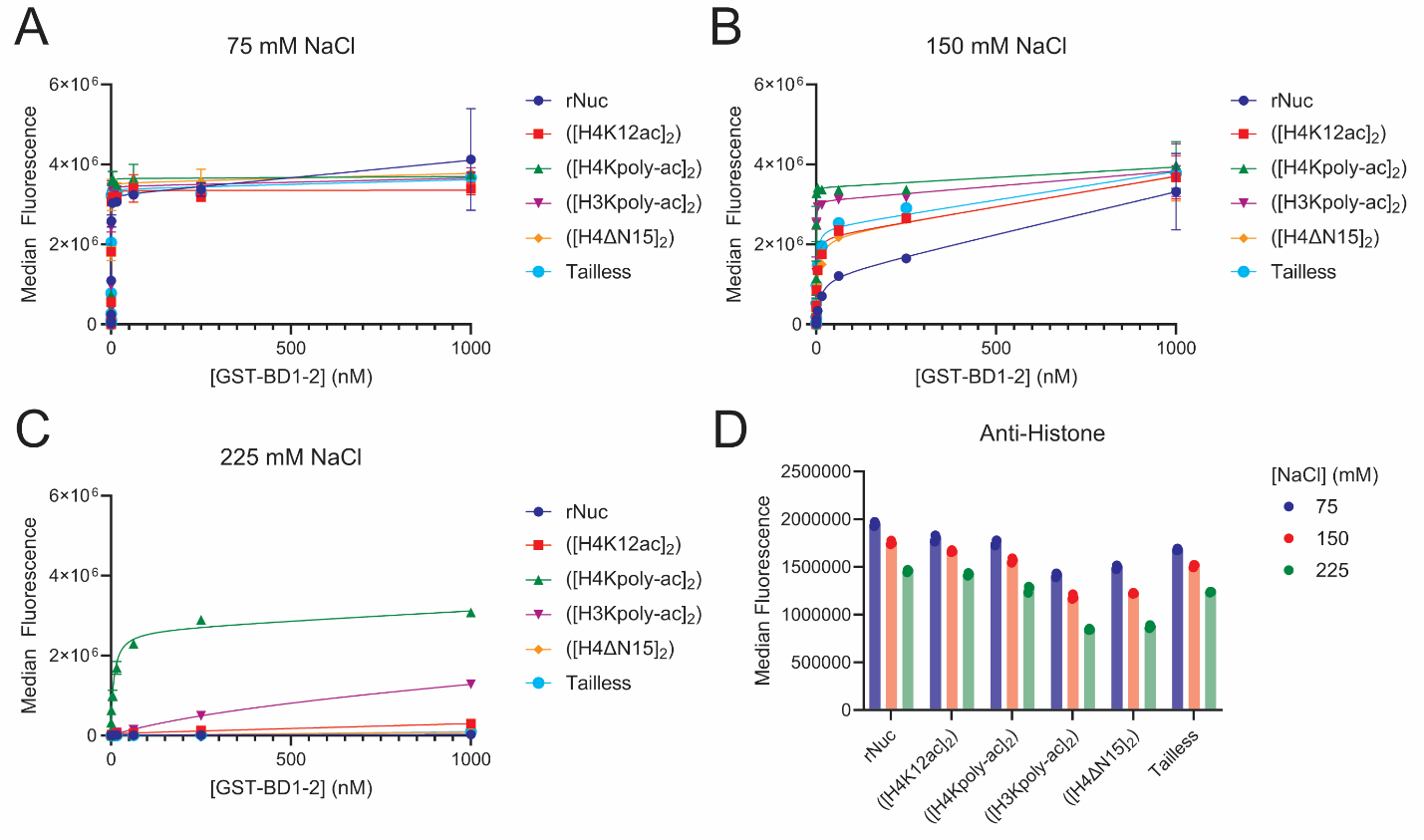

**Supplemental Figure S3.** GST-BRD4 BD1-2 binding to nucleosomes in Luminex binding assays in different NaCl concentration buffers. The nucleosome panel included unmodified recombinant nucleosome, H4K12 acetylated ([H4K12ac]_2_), H4 poly-acetylated ([H4K5acK8acK12acK16ac]_2_), H3 poly-acetylated ([H3K4acK9acK14acK18ac]_2_), H4 tail truncated ([H4ΔN15]_2_), and trypsin digested tailless nucleosomes (see Methods for bead coupling and the Luminex assay). (**A-C**) GST-BD1-2 nucleosome binding titrations using buffer containing (**A**) 75 mM NaCl, (**B**) 150 mM NaCl, and (**C**) 225 mM NaCl. Assays were repeated in triplicate and binding curves were fitted to a one-site binding model in GraphPad Prism. (D) Control Luminex assays detecting total histone levels using a general histone antibody in each of the three buffer conditions tested. Titrations and controls were run in triplicate.

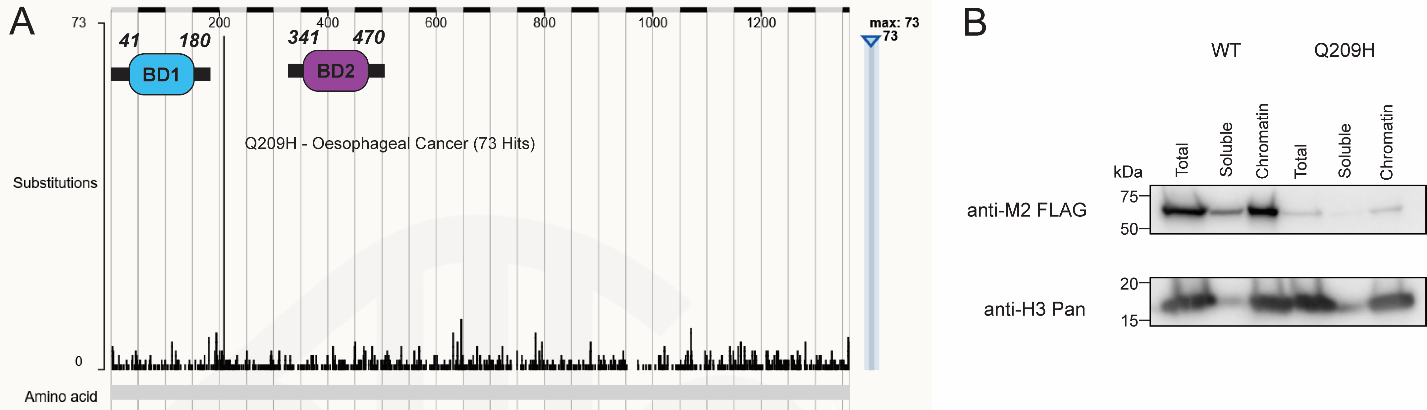

**Supplemental Figure S4.** Cancer-linked mutation (Q209H) in the BRD4 linker region does not affect chromatin binding. (**A**) COSMIC database illustration showing frequency of cancer-associated mutations in human BRD4, highlighting the frequently occurring Q209H mutation. (**B**) Chromatin association assays comparing wild-type BRD4 BD1-2 wild-type and Q209H mutant in HEK293T cell extracts. Experiments were performed in biological triplicates.

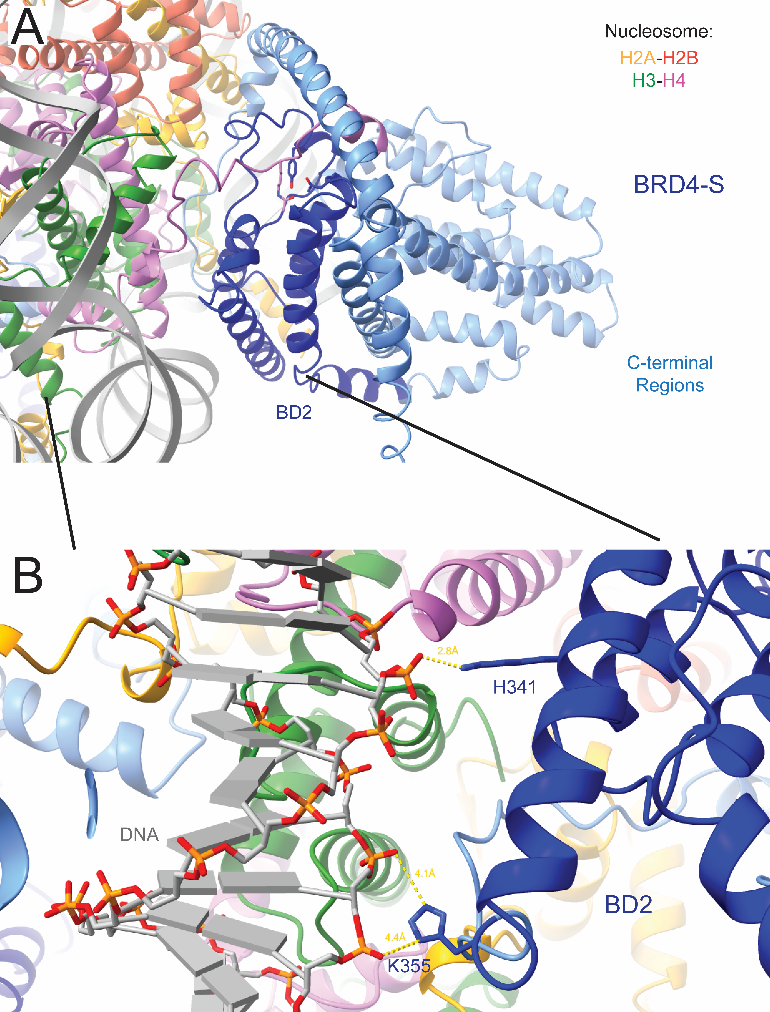

**Supplemental Figure S5.** Potential BD2 interactions with DNA based on AlphaFold model of BRD4-S bound to [H4K12ac]_2_ nucleosome. (**A**) View of the BRD4-S BD2 engaging [H4K12ac]_2_ and nucleosomal DNA. (**B**) Zoomed in view of the BD2 highlighting basic residues H341 and K355 that potentially interact with the nucleosomal DNA phosphodiester backbone.

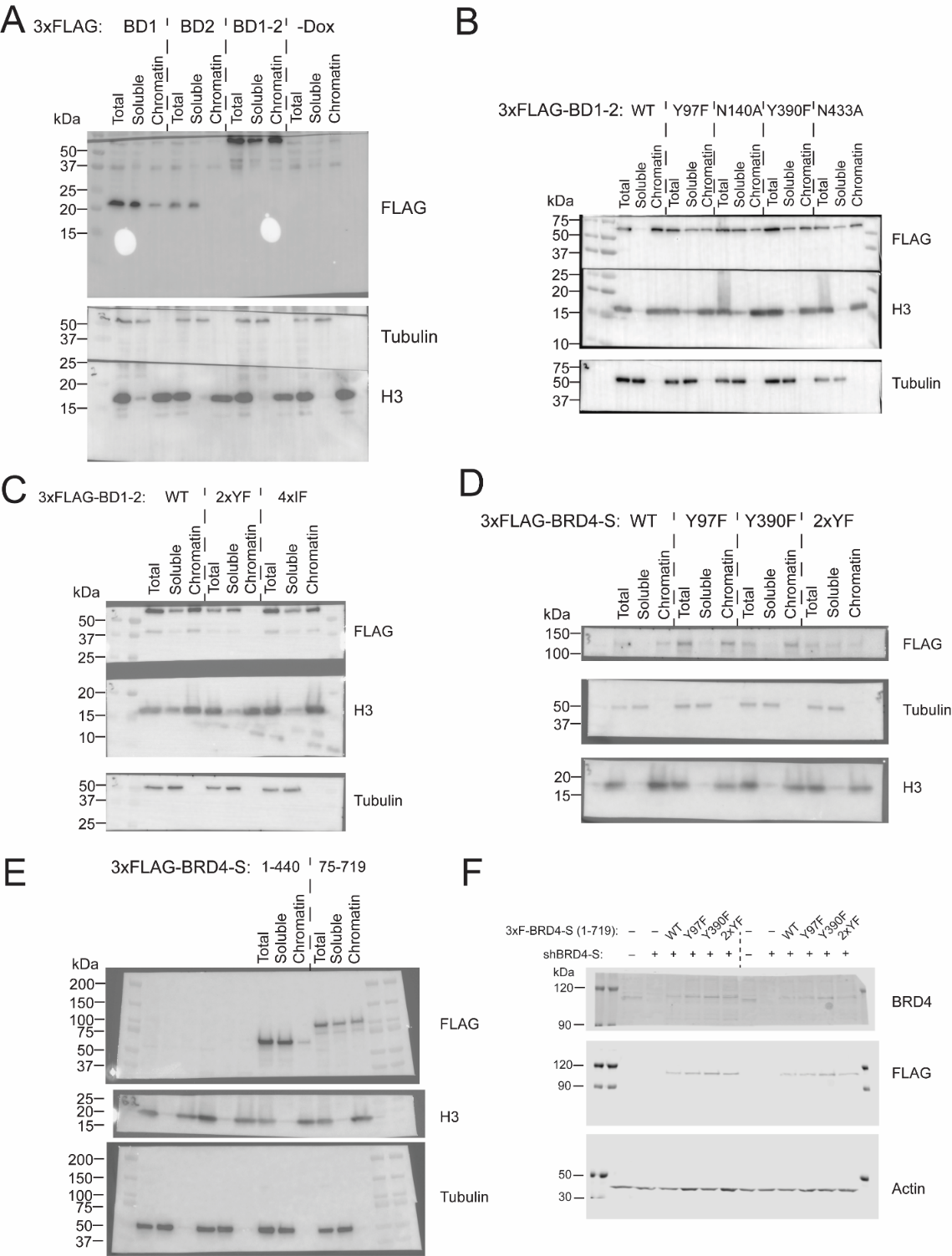

**Supplemental Figure S6.** Western blots of BRD4 chromatin association assays. (**A**) Western blots for BRD4 BD1, BD2, and BD1-2 truncation series chromatin association assays. Samples were rerun and blotted for control band clarity. (**B**) Western blots BRD4 BD1-2 pocket mutant chromatin association assays. (**C**) Western blots for BRD4 BD1-2 double pocket mutant (2xYF) and four putative interface mutant (4xIF) chromatin association assays. (**D**) Western blots for BRD4-S mutant chromatin association assays. (**E**) Western blots for BRD4-S truncation 2xYF mutant chromatin association assays. (**F**) Western blots for BRD4-S mutant MDA-MB-231 cancer biology experiments. All chromatin association assays were run in biological triplicate for band quantifications.
